# Segment Any Plant (SAP): Foundation-Model Segmentation for Plant Time-Series Phenotyping

**DOI:** 10.64898/2026.03.11.711099

**Authors:** Alex Abbey, Yasmine Meroz

## Abstract

Quantitative studies of plant growth and environmental responses increasingly rely on time-series imaging, yet automated segmentation remains challenging due to continuous growth, large non-rigid morphological change, and frequent self-occlusion. Traditional image-processing pipelines and task-specific deep learning models often require extensive annotated datasets and retraining, limiting portability across species, developmental stages, and imaging conditions. Here we present SAP (Segment Any Plant), a plant-focused framework that leverages the pretrained Segment Anything Model 2 (SAM2) to enable few-shot, training-free segmentation of plant time-series imagery. SAP integrates interactive prompting, automated temporal mask propagation, and centerline extraction within a web-based interface, allowing users to move from raw images to quantitative descriptors of organ shape and dynamics without programming expertise. Across multiple systems, including Arabidopsis thaliana rosette development, root growth, sunflower gravitropism, and confocal root microscopy, SAP achieves high segmentation accuracy (mean IoU 0.89–0.93) and sub-pixel centerline precision from single-frame prompting. By reducing the need for task-specific retraining, SAP provides a transferable framework for reproducible time-series phenotyping across diverse experimental contexts.

## Introduction

Time-resolved imaging has become central to quantitative plant biology, enabling measurement of growth, development, and environmental responses across laboratory and field settings. Because plant form is inherently dynamic, its quantitative characterization increasingly depends on long time-series imaging across spatial scales, from cells to whole organs. Unlike many other biological systems, plants continuously grow and undergo large, non-rigid morphological changes. This intrinsic plasticity, often accompanied by self-occlusion, branching, and topological change, presents a fundamental challenge for automated image analysis. Segmentation and tracking pipelines that perform well for static or shape-constrained systems frequently degrade when applied to extended plant time series, limiting their robustness and generality.

Early phenotyping approaches relied on rule-based image processing, including thresholding, morphological operations, and geometric feature extraction. Tools such as PlantCV (1) and specialized root and shoot pipelines (2–5) provide efficient and interpretable solutions but are sensitive to background complexity, illumination variability, and organ overlap. Deep learning methods have improved segmentation accuracy across roots, leaves, shoots, and meristems (6– 10), yet typically require large annotated datasets and task-specific training. In practice, each new species, imaging modality, or developmental context often necessitates retraining, creating a recurring bottleneck between image acquisition and quantitative analysis. As imaging throughput increases, the limiting factor in plant phenotyping is no longer data collection, but the scalability and adaptability of segmentation workflows.

Recent advances in machine learning have introduced foundation models: large-scale vision systems trained on diverse datasets that learn transferable visual representations. Rather than optimizing a model for a single predefined task, foundation models enable prompt-based adaptation with minimal supervision. In computer vision, the Segment Anything Model 2 (SAM2) enables few-shot, interactive segmentation in images and videos without task-specific retraining (11). This shift from bespoke supervised models to general-purpose visual priors represents a conceptual change in how segmentation problems can be approached. Despite their promise, foundation models have seen limited application in plant phenotyping (12). Plants pose a stringent test case: they exhibit continuous growth, large deformations, organ emergence, and self-occlusion across extended time-series experiments. Whether foundation models can accommodate these features without retraining remains largely unexplored. If successful, such an approach would reduce annotation burdens, increase methodological portability across species, and accelerate quantitative analysis in new experimental systems.

Here, we introduce SAP (Segment Any Plant), a plant-focused framework that leverages SAM2 to enable rapid, few-shot segmentation and analysis of plant time-series imagery. Rather than developing a task-specific model, we demonstrate that a general-purpose foundation model can be transformed into a robust plant segmentation and tracking pipeline through minimal interactive prompting and temporal propagation. SAP integrates a web-based interface with automated mask propagation and centerline extraction, allowing researchers to move from raw images to quantitative descriptors of organ growth and morphodynamics without programming expertise.

We validate SAP across diverse systems, including Arabidopsis thaliana rosette development, root growth, sunflower gravitropism, and confocal root microscopy. Across biological scales, SAP maintains high segmentation accuracy from single-frame prompting while preserving temporal stability across extended sequences. Beyond performance benchmarking, we show that segmentation and centerline extraction enable quantitative analyses of growth rates, curvature dynamics, and organ-level morphodynamics.

By aligning foundation-model segmentation with the specific challenges of plant growth, SAP provides a scalable framework for time-series phenotyping. More broadly, it illustrates how interactive foundation models may serve as a new computational infrastructure for plant science, lowering technical barriers and expanding the range of experimentally accessible questions in development and environmental response.

## Methods

### Computational Requirements

The computational requirements for running SAP are primarily determined by the underlying SAM2 model. For inference, GPU memory requirements range from 8GB VRAM for the smaller model variants (tiny and small) processing standard resolution imagery, to 12-24GB for the larger models or higher resolution inputs. While SAM2 was originally trained on 256 NVIDIA A100 GPUs with 80GB memory each (11), inference on modern consumer-grade GPUs (such as RTX 4060 or higher) is feasible for typical plant phenotyping applications. Real-time processing performance varies with hardware: the large model achieves approximately 30 frames per second on an A100 GPU, while smaller variants can process around 47 frames per second on the same hardware. This scalability allows researchers to select model-hardware configurations appropriate to their computational budget and throughput requirements.

### Datasets Descriptions

Five datasets spanning different species, organs, imaging modalities, and biological scales were used to evaluate SAP (Table 1).

**Table 1.** Summary of datasets and their role in validation. Checkmarks indicate availability of ground-truth annotations for mask and centerline validation, respectively. Each dataset is accompanied by a supplementary video.

| Dataset | Modality | Mask GT | Centerline GT | Application |
| --- | --- | --- | --- | --- |
| *Arabidopsis* rosette (9) (Video 2) | Top-view time-lapse | ✓ | – | Leaf growth tracking |
| Sunflower gravitropism (Video 3) | Side-view time-lapse | ✓ | ✓ | Gravitropic response |
| Confocal root cells (13) (Video 5) | Confocal *z*-stack | ✓ | – | Cell-level *z*-tracking |
| *Arabidopsis* root growth (Video 4) | Side-view time-lapse | – | ✓ | Root elongation |
| Confocal root time-lapse (14) (Video 6) | Confocal time-lapse | – | – | Live cell tracking |

#### High-Throughput phenotyping of Arabidopsis thaliana rosette growth

The dataset (9) consists of high-throughput plant phenotyping image data acquired from *Arabidopsis thaliana* plant trays. The images used for validation were taken once a day for 9 consecutive days, each of one of 64 plants, and were hand-tagged. The dataset thus includes 576 ground-truth hand-tagged masks. We joined the whole dataset into a single video file, essentially showing the growth of multiple plants over time, and tagged one frame. SAP then segmented the plant in the tagged frame, and the mask was propagated through the entire video sequence (Supplementary Video 2).

#### Sunflower gravitropism dataset

This dataset is based on unpublished experiments we ran for testing gravitropism in sunflower (*Helianthus annuus*) plants. The images used for validation are images of 3 defoliated sunflowers, taken every 15 minutes for 32 hours, and were hand-tagged. The dataset thus includes 384 ground-truth hand-tagged masks and centerlines (Supplementary Video 3).

#### Confocal microscopy of Arabidopsis root cells

To evaluate SAP on sub-cellular microscopy data, we used confocal *z*-stack imagery of *Arabidopsis thaliana* roots from the dataset of Strauss et al. (13). A 51-slice substack of the Propidium Iodide channel was selected, spanning the *z*-range of individual cells. Ground-truth mask segmentation was obtained from a MorphoGraphX watershed label volume distributed with the dataset. Three individual cells were selected for quantitative comparison (Supplementary Video 5).

#### Arabidopsis root growth dataset

This dataset is based on unpublished experiments we ran for testing growth and memory in *Arabidopsis* plants. The plants were grown in petri dishes in the dark and photos were taken with a dim green light. Images were taken every 9 minutes for 30 hours, creating 200 images altogether. Ground-truth centerlines were hand-tagged for this dataset (Supplementary Video 4).

#### Confocal time-lapse of Arabidopsis root growth

To demonstrate SAP at the cellular scale on live imaging data, we used a confocal time-lapse of a vertically growing *Arabidopsis thaliana* root from the dataset of von Wangenheim et al. (14). No ground-truth segmentation was available for this dataset; it is included as a qualitative demonstration of cell-level tracking (Supplementary Video 6).

### Quantitative Performance Metrics

Segmentation accuracy was assessed using the Intersection over Union (IoU) metric, which is commonly used to measure the performance of object category segmentation methods(15). The IoU is calculated for each image pair as:

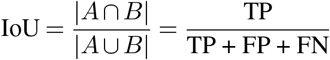

where *A* represents the predicted segmentation mask, *B* denotes the ground truth annotation, and TP, FP, and FN correspond to true positive, false positive, and false negative pixel classifications, respectively. IoU= 1 reflects perfect overlap between prediction and ground truth, while IoU= 0 indicates no overlap; the metric penalizes both missed regions (under-segmentation) and spurious regions (oversegmentation), making it a stringent measure of boundary accuracy.

Descriptive statistics were computed across all samples, including mean IoU 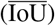, standard deviation (*σ*), and confidence intervals.

To evaluate model consistency throughout time, Pearson correlation analysis was performed between temporal progression and segmentation performance. The correlation coefficient was computed as:

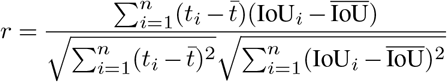

where *t*_*i*_ represents discrete time points. A low correlation means IoU values are stable over time: performance does not drift as plants move, bend, or change morphology.

## Results

### Software Implementation

SAP is a web application built on top of Meta’s open-source SAM2 interactive demo (11). It follows a client–server architecture with three main components: a browser-based frontend for user interaction and visualization, an inference backend that hosts the SAM2 model and handles segmentation and mask propagation, and a video processing backend responsible for frame extraction and plant-specific analyses such as centerline and skeleton extraction.

Users interact with the application through a point-and-click interface in the browser. Segmentation masks are computed on the server, streamed back to the client, and displayed as overlays on the video frames in real time. The video processing backend uses standard scientific Python libraries (OpenCV, scikit-image, SciPy) for morphological operations and centerline computation.

SAP supports four SAM2.1 model variants (tiny, small, base_plus, and large) paired with three resolution presets (1024, 1536, or 2048 px), allowing users to balance processing speed and segmentation quality for their available hardware. Inference runs on CUDA-enabled GPUs, with fallback support for Apple Silicon and CPU. The application is deployed via Docker and is available as an open-source repository (https://github.com/merozlab/plant-segmentation-app).

### Prompting and Segmentation Workflow

In this section, we provide a step-by-step description of the segmentation workflow implemented in the SAP interface, which is also illustrated in Fig. 1. The workflow guides the user from initial data preparation through segmentation, temporal propagation, manual corrections, and result extraction.

**Fig 1.**
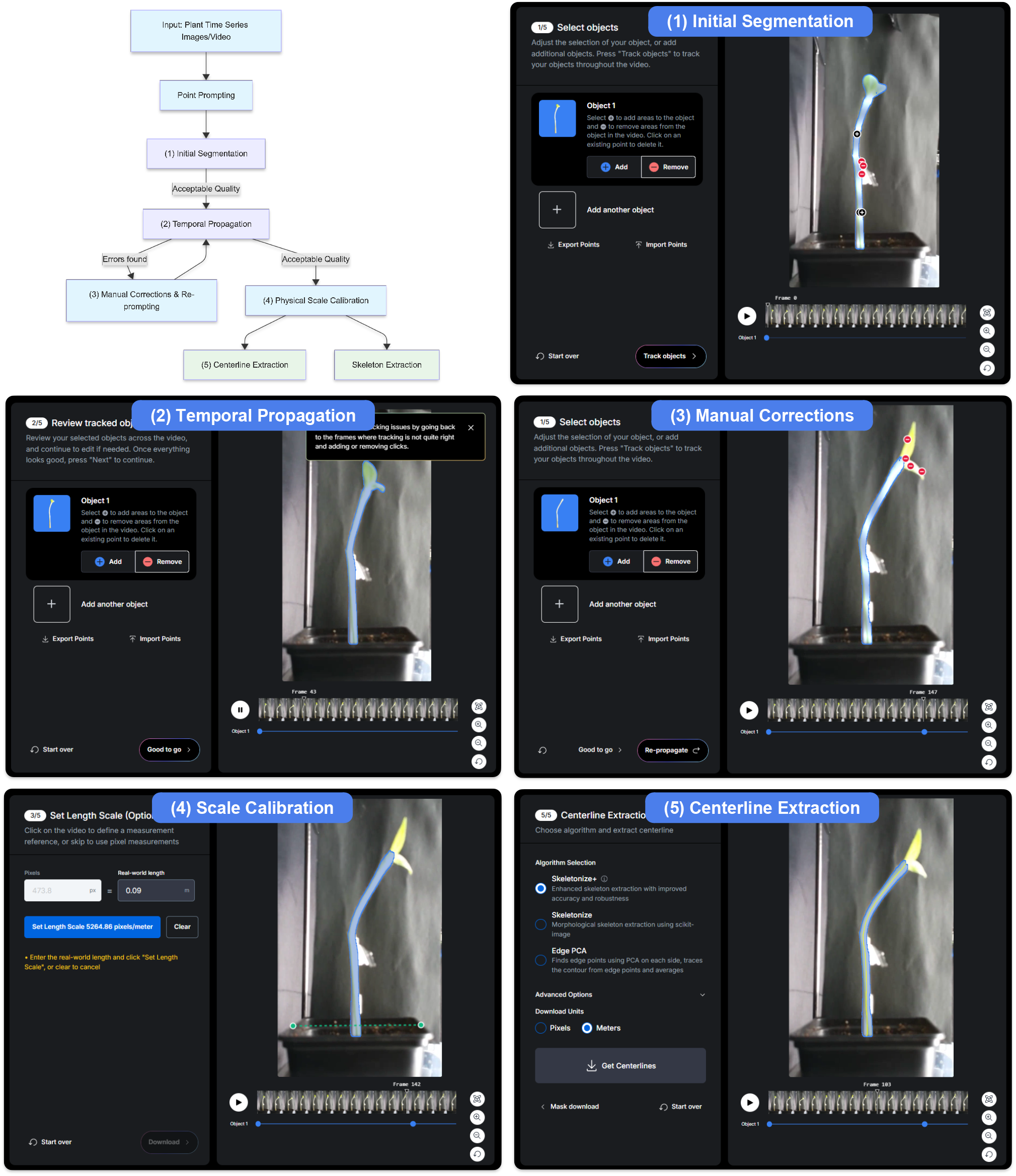
Overview of the SAP workflow. Top left: flowchart of the processing pipeline. Top right and below: corresponding screenshots from the SAP interface. (1) **Initial Segmentation**: the user provides point prompts on the plant of interest to generate segmentation masks on selected frames. (2) **Temporal Propagation**: masks are automatically propagated across the full image sequence. (3) **Manual Corrections**: if propagation quality is insufficient, the user refines masks with additional prompts and re-propagates. (4) **Scale Calibration**: a known reference length is used to convert pixel measurements to real-world units. (5) **Centerline Extraction**: a centerline or skeleton is extracted from the segmentation masks for morphological analysis. See Methods for a full description of each step. A video demonstration of this workflow is provided in Supplementary Video S1.

#### Preprocessing: Data Upload and Cropping

The user begins by uploading a video file or a sequence of images. Optionally, the video can be cropped to focus on a region of interest, which is particularly useful for large datasets or sequences containing multiple plants. Cropping also helps accommodate SAM2’s maximum resolution limits (see Discussion for details).

#### Step 1: Initial Segmentation

Initial frames are selected for segmentation, where the user provides point prompts on the plant of interest. SAP generates a segmentation mask for these frames. The interface supports multiple plants within a frame, with the maximum number configurable depending on available computational resources.

#### Step 2: Temporal Propagation

The initial masks serve as a seed for the rest of the frames. Using SAM2’s mask propagation, the segmentation is automatically transferred across the sequence, enabling tracking of morphological changes and positional shifts over time. Propagation can run forwards and backwards from an annotated frame, ensuring full coverage even when plants emerge or change significantly.

#### Step 3: Manual Corrections

For frames where automatic propagation is insufficient, users can refine the mask with additional positive or negative prompts. This iterative process maximizes accuracy on challenging frames while minimizing manual effort on well-segmented ones. Segmentation masks are overlaid on the original frames for quick quality verification.

#### Step 4: Scale Calibration

Once segmentation is complete, the user defines a length scale by selecting a known reference distance within the video. This enables conversion from pixel measurements to real-world units.

#### Step 5: Centerline Extraction

Finally, the interface allows extraction of a centerline or skeleton from the segmentation mask. This is particularly useful for analyzing elongated structures such as stems or roots. Centerlines are immediately visualized in the interface for quality assessment and are also available for export. Details of the extraction method are provided in Appendix A.

### Mask Validation

Table 1 summarizes all five datasets used in this study and indicates which validation task each supports. Four datasets include ground-truth annotations: three with hand-tagged masks for segmentation validation, and two with hand-tagged centerlines for centerline validation (the sunflower dataset has both). The remaining dataset serves as a qualitative demonstration. We compared SAP-predicted masks against ground truth across the three mask-labeled datasets (Table 2). The first two temporal datasets required only a single initial frame annotation, from which masks were automatically propagated through the sequence.

**Table 2.** IoU and temporal correlation statistics for SAP vs. ground-truth masks. *r* = Pearson(*t*, IoU) is the temporal correlation coefficient, computed per object; |*r*| ≈ 0 indicates stable segmentation across frames. Frames where both SAP and GT masks are absent (cell outside the optical section) are excluded from all confocal statistics.

| Metric | *Arabidopsis* growth | Sunflower grav. | Confocal root |
| --- | --- | --- | --- |
| Sample size $N$ | 576 | 384 | 121 |
| Objects tracked | 64 | 48 | 3 |
| Mean IoU | $0.886 \pm 0.042$ | $0.927 \pm 0.016$ | $0.75 \pm 0.19$ |
| Mean $r$ | $0.44 \pm 0.35$ | $0.19 \pm 0.10$ | – |

For the *Arabidopsis* rosette dataset (9) (Video 2), the automated method achieved a mean IoU of 0.88 ± 0.042 across all 576 frames. The sunflower gravitropism dataset (Video 3) showed higher segmentation accuracy, with a mean IoU of 0.927 ± 0.016 across 384 frames. These IoU values demonstrate strong overlap between predicted and ground truth masks, indicating robust segmentation performance across both datasets. The confocal root dataset (13) (Video 5) yielded per-cell mean IoU values of 0.809 ± 0.018 (*n* = 51), 0.65 ± 0.29 (*n* = 36), and 0.77 ± 0.16 (*n* = 34), with the reduced sample sizes for Cells 2 and 3 reflecting slices where the optical section exits the tissue and both SAP and GT masks vanish (Fig. 2D).

**Fig 2.**
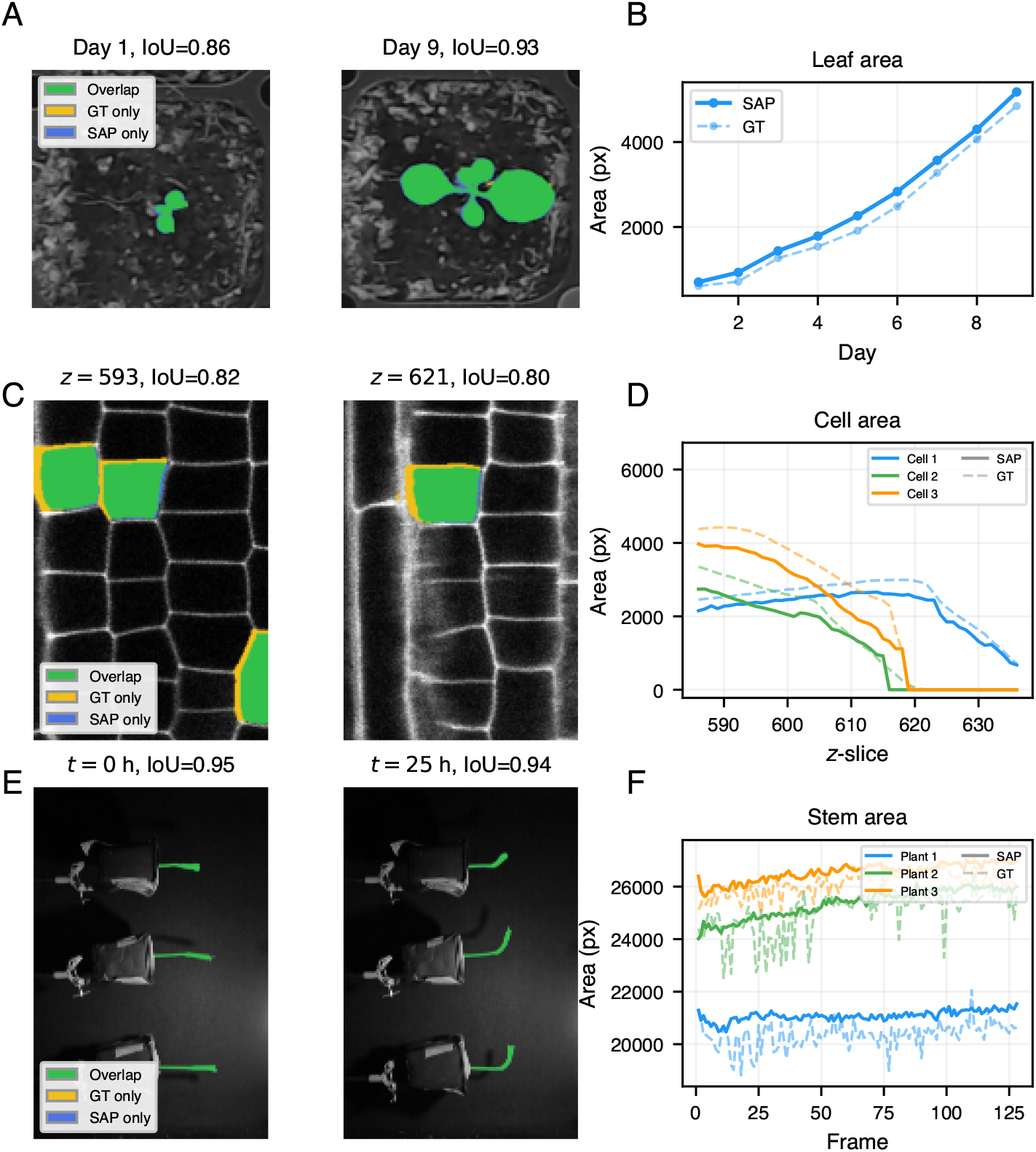
Validation of SAP segmentation against manual ground truth. Top row: *Arabidopsis thaliana* rosette growth (Lee et al. (9); Video 2). (A) Overlay of SAP masks and ground truth at Day 1 (IoU = 0.86) and Day 9 (IoU = 0.93); green: overlap, orange: ground truth only, blue: SAP only. (B) Leaf area tracked by SAP and ground truth over 9 days of growth. Middle row: confocal *Arabidopsis* root cells (Strauss et al. (13); Video 5). (C) Overlay of SAP and ground-truth masks for three cells at *z* = 593 (IoU = 0.82) and *z* = 621 (IoU = 0.80). (D) Cross-sectional area tracked by SAP (solid) and ground truth (dashed) for each cell across *z*-slices. Bottom row: sunflower (*Helianthus annuus*) gravitropism (Video 3). (E) Overlay of SAP and ground-truth masks for three plants at *t* = 0 h (IoU = 0.95) and *t* = 25 h (IoU = 0.94). (F) Stem area tracked by SAP (solid) and ground truth (dashed) for each of the three plants over 128 frames (32 h). In all three datasets, SAP accurately captures object boundaries and faithfully tracks area changes over time and space.

Temporal stability was assessed using Pearson correlation coefficients between IoU values and frame time. The sunflower gravitropism dataset exhibited low temporal correlation (mean *r* = 0.19 ± 0.10), indicating that segmentation accuracy remained stable despite continuous changes in plant orientation during the gravitropic response. The *Arabidopsis* growth dataset showed moderate temporal correlation (mean *r* = 0.44 ± 0.35), likely reflecting systematic changes in segmentation accuracy as plants underwent substantial morphological development over the nine-day period, including leaf emergence and expansion. For the confocal root cells, the correlation between SAP and GT cross-sectional areas were low ( − 0.17, *p* = 0.22) for the cell that existed throughout all frames, yet very negative for ( − 0.66, *p* = 1.4 × 10^−5^; − 0.74, *p* = 5.9 × 10^−7^) for cells that disappeared during the z-stack, indicating that SAP did not accurately capture the disappearance of cells as they exit the optical section (Fig. 2D).

These validation metrics demonstrate that SAP achieves high segmentation accuracy across diverse plant imaging scenarios with minimal user input, maintaining consistent performance throughout temporal sequences despite significant morphological and positional changes.

### Centerline Extraction Validation

To validate centerline extraction, we compared automated centerlines against ground truth on two datasets: 128 frames from the sunflower gravitropism dataset (Video 3) and 200 frames from the *Arabidopsis* root growth dataset (Video 4) (Table 3, Fig. 3B, E). Ground truth centerlines were generated by manually tagging start and end points, then applying the same contouraveraging algorithm used in the automated method.

**Table 3.** Centerline extraction accuracy. Point-wise Euclidean distances and endpoint localization errors between SAP-extracted and manually annotated ground truth centerlines for both centerline-validated datasets (Table 1). Both centerlines were interpolated to 256 equidistant points before comparison. Values are reported as mean *±* standard deviation.

| Metric | Sunf. grav. | Arab. root |
| --- | --- | --- |
| Sample size $N$ | 128 frames | 200 frames |
| Mean Distance Error (px) | $1.06 \pm 0.78$ | $1.03 \pm 0.32$ |
| RMSE (px) | $1.23 \pm 0.88$ | $1.17 \pm 0.36$ |
| Mean Endpoint Error (px) | $2.3 \pm 1.0$ | $2.1 \pm 0.5$ |

**Fig 3.**
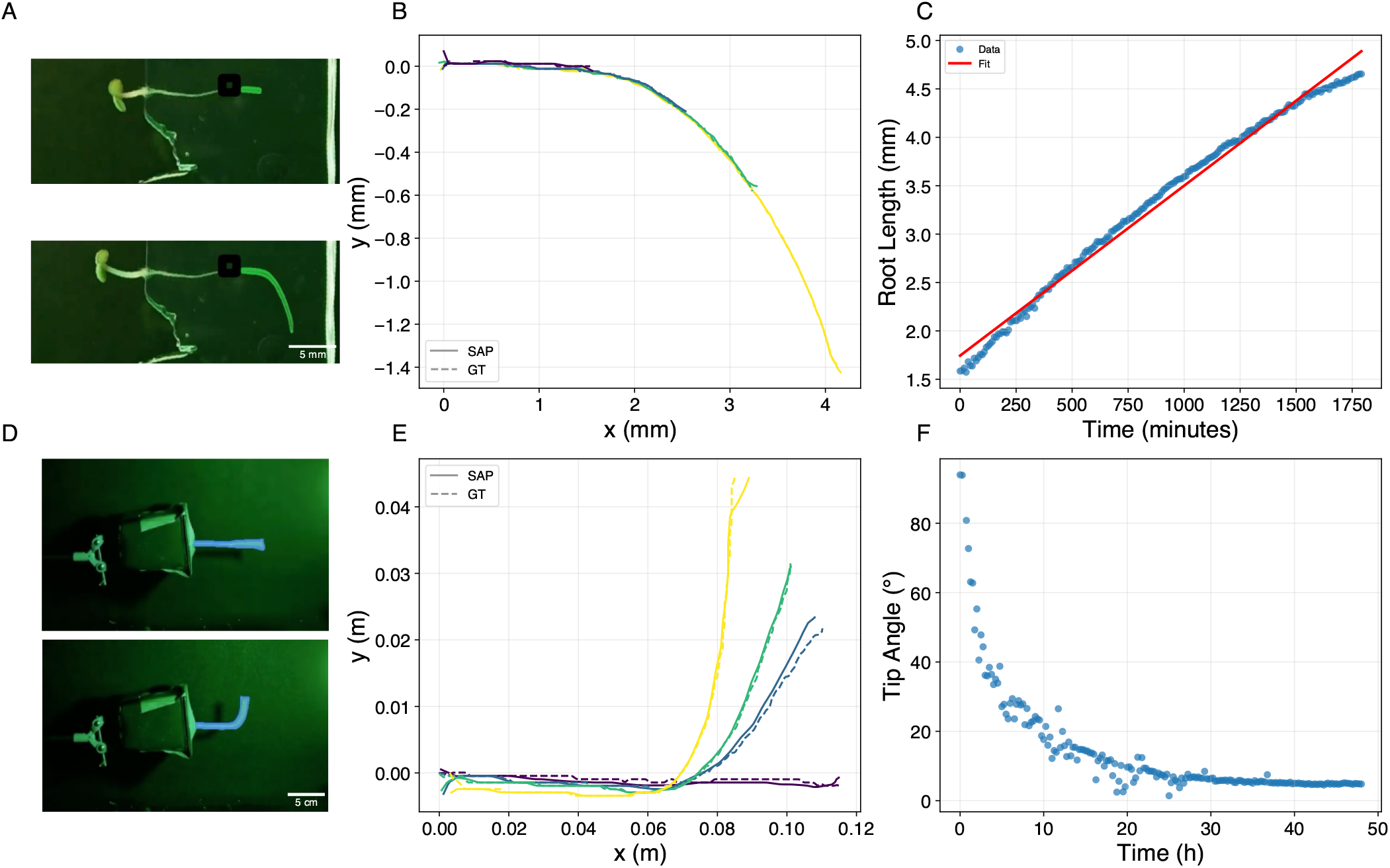
Quantitative analyses of plant organ dynamics with SAP. Top row: *Arabidopsis thaliana* root growth (Video 4). (A) Representative frames showing SAP segmentation masks and extracted centerlines at an early and late time point. (B) Root centerlines over ∼ 30 h, colored from yellow (early) to purple (late); SAP-extracted (solid) and manually annotated ground truth (dashed) centerlines are overlaid, showing close agreement. (C) Root length versus time fitted by a linear model (*R*^2^ = 0.99), with elongation rate 1.76 *×* 10^−3^ mm min^−1^ . Bottom row: *Helianthus annuus* (sunflower) gravitropism (Video 3). (D) Representative frames showing SAP segmentation masks and extracted centerlines at an early and late time point. (E) Stem centerlines during gravitropism, colored from yellow (early) to blue (late); SAP-extracted (solid) and ground truth (dashed) are overlaid. (F) Tip angle versus time, showing the gravitropic response over ∼50 h.

For each frame, point-wise Euclidean distances were computed between corresponding points on the automated and ground truth centerlines (both interpolated to 256 points). For the sunflower dataset, the mean distance error was 1.06 ± 0.78 pixels (RMSE: 1.23 ± 0.88 pixels) with endpoint error of 2.3 ± 1.0 pixels. For the *Arabidopsis* root dataset, comparable accuracy was achieved with a mean distance error of 1.03 ± 0.32 pixels (RMSE: 1.17 ± 0.36 pixels) and endpoint error of 2.1 ± 0.5 pixels. In both cases, mean errors represent approximately one pixel, confirming sub-pixel centerline accuracy suitable for quantitative analysis of organ shape and growth dynamics.

### Time-Series Analysis Demonstrations

The preceding sections validated SAP’s mask and centerline accuracy on the four ground-truth datasets (Table 1). Here we highlight quantitative analyses enabled by centerline extraction, as well as a qualitative demonstration on the confocal root time-lapse dataset, which lacks ground-truth annotations.

The sunflower gravitropism dataset (Video 3) consists of three plants in the same frame (only one shown in Fig. 2), demonstrating SAP successfully tracking multiple objects simultaneously. The *Arabidopsis* rosette growth dataset (Video 2) illustrates SAP’s ability to cope with substantial morphological changes over a 9-day period, including the emergence of new leaves. Figure 3 shows quantitative analyses that centerline extraction enables on the *Arabidopsis* root growth dataset (Video 4) and the sunflower dataset: fitting growth models to root elongation (using centerlines) (19), and extracting tropism parameters using the AC model (20). To further demonstrate SAP’s versatility at the cellular scale, we applied it to the confocal root time-lapse dataset (14) (Table 1). By tagging a few cells in a single confocal slice, SAP propagated the segmentation across subsequent frames, tracking individual cells as the root elongated. Although no ground truth was available, the resulting masks visually follow cell boundaries throughout the sequence, suggesting that SAP can be extended to cell-level tracking in live confocal microscopy (Supplementary Video 6). Representative frames from each dataset are shown in Supplementary Figs. 2–6. We note that thoughtful prompting creates better results and saves time during the segmentation process. For example, in the sunflower gravitropism dataset, we prompted SAP with a point on each plant in the first frame, and it was able to segment all three plants in the subsequent frames without further user input. In the growth experiment, segmentation worked much better when prompting SAP with a point on a fully grown plant, as SAP will propagate backwards and achieves higher accuracy for the earlier frames.

## Discussion

SAP makes the capabilities of SAM2 accessible to plant scientists through an interactive interface that requires no coding expertise and integrates mask propagation with centerline extraction. Our results show that foundation models can be adapted into practical tools for plant phenotyping, accommodating growth-driven deformation and temporal dynamics without task-specific retraining. By coupling pretrained visual representations with plant-focused geometric analysis, SAP provides a flexible framework for time-series image analysis across diverse experimental systems.

Supervised deep-learning approaches have achieved high segmentation accuracy in many plant imaging contexts, often reporting mIoU values above 0.8 on in-domain datasets (8, 16). However, these methods typically require extensive annotated training data and model customization. In contrast, SAP achieves IoU values of 0.89 for Arabidopsis and 0.93 for sunflower (Table 2) using single-frame prompting and temporal propagation (Table 4). While comparable in performance, the principal advantage lies in portability: segmentation can be deployed across species, organs, and imaging modalities without retraining. This reduction in annotation burden is particularly relevant for exploratory experiments, small-scale studies, or systems where annotated datasets are limited.

**Table 4.** Comparison of plant segmentation approaches. SAP is compared with classical thresholding-based methods (KymoRod) and supervised deep learning models (U-Net/DeepLabV3+) across training requirements, user input, occlusion handling, and key limitations.

| Feature | SAP | KymoRod (5) | U-Net / DeepLabV3+ (e.g., (8, 16)) |
| --- | --- | --- | --- |
| **Training Requirements** | No training (uses SAM2 foundation model) | No training (automatic or manual thresholding) | Extensive task-specific training data required |
| **User Input** | Minimal – annotate initial frame only | Threshold tuning; may require manual organ separation in ImageJ | Task- and plant-specific annotations |
| **Approach** | Foundation model; versatile across diverse plants and organs | Thresholding-based contour detection | Deep learning with task-specific models |
| **Occlusion Handling** | Handles occlusions | Cannot handle occlusions | Depends on model and training data |
| **Lighting Conditions** | Works in various conditions | Requires controlled conditions and high contrast | Standard lighting preferred |
| **Key Limitations** | Max resolution 2048px; requires initial annotation | Requires Matlab; sensitive to background and contrast | Resource-intensive; requires technical expertise; limited portability across species |

Classical threshold-based methods such as KymoRod (5) avoid supervised training but are sensitive to contrast conditions and struggle with occlusion. Foundation-model segmentation offers an alternative strategy that combines largescale visual priors with minimal user guidance. In the timeseries datasets examined here, propagation from a single annotated frame maintained stable performance despite sub-stantial morphological change. Such temporal consistency is essential for morphodynamic studies, where small segmentation drifts can accumulate over long sequences and bias downstream quantitative measurements.

Beyond segmentation, SAP incorporates a contour-averaging centerline extraction algorithm tailored to elongated plant organs. Traditional thinning-based skeletonization (18) may deviate toward corners at organ tips, and alternative approaches such as Voronoi skeletonization (5) or intensity-based marching cross-sections (3) depend on high contrast or per-frame manual input. By separating mask generation from geometric analysis, SAP adopts a modular strategy in which robust segmentation is followed by deterministic contour-based centerline construction. This approach yields sub-pixel centerline accuracy (Table 3) and enables integration of exported centerlines with downstream kinematic, biomechanical, or growth modeling analyses (Table 5).

**Table 5.** Comparison of centerline and skeleton extraction methods. SAP’s contour-averaging approach is compared with existing methods for extracting organ midlines, including Voronoi skeletonization (KymoRod), marching cross-sections (Interekt), trained skeleton prediction (SLEAP), contour curvature analysis, and morphological thinning. Columns compare required input, algorithmic approach, endpoint accuracy, and key limitations.

| Feature | SAP | KymoRod (5) | Interekt (3) | SLEAP (4) | Yamamoto & Haga (17) | Morphological Thinning (e.g., (18)) |
| --- | --- | --- | --- | --- | --- | --- |
| **Input Required** | Segmentation mask | Thresholded contour | Raw image & base point | Raw image | Organ contour & manually seperated edges | Segmentation mask |
| **Method** | Find tips and connect via contour averaging | Voronoi skele-tonization of contour | Marching cross-section; midpoint of intensity edges | Trained skeleton prediction | Curvature analysis along contour | Mathematical thinning algorithms |
| **Tip/Endpoint Accuracy** | Accurate at tips via contour averaging | May deviate at tips (skeleton branches) | Depends on edge detection quality | Varies with training data | Not specified | Deviates toward corners at tips |
| **Key Limitations** | Requires segmentation mask as input | Requires Matlab; needs high contrast and surface texture for PIV | Manual input per frame; no occlusion handling | High user input for training; less flexible than mask-based | Requires manual edge separation; limited to simple geometries | May not reach edges accurately; points toward corners |

SAP’s adaptive resolution presets further support use across heterogeneous imaging contexts, from confocal microscopy to whole-plant phenotyping platforms. Although initial prompting is required, the few-shot workflow substantially reduces manual effort relative to frame-by-frame annotation. Fully automated deployment, for example in high-throughput or field-based imaging, may benefit from integration with automated object detection in future implementations.

As with any segmentation method, performance depends on image quality and experimental context. Extremely low-contrast images or highly complex canopies may reduce segmentation fidelity, and continued evaluation across additional plant architectures and environmental conditions will further define the operational range of the method. Importantly, the modular design of SAP, in which foundation-model segmentation is coupled to plant-specific geometric analysis, allows adaptation and refinement as new use cases emerge. Rather than replacing domain-specific expertise, foundation models provide a flexible computational basis upon which plant-focused analytical tools can be constructed, combining general visual representations with biologically informed post-processing.

In summary, SAP demonstrates that foundation-model segmentation can support robust analysis of plant growth and morphodynamics across biological scales. By combining general-purpose visual representations with plant-specific geometric postprocessing, SAP provides a scalable and adaptable approach to time-series phenotyping that lowers technical barriers while preserving quantitative rigor. As imaging experiments increasingly span extended developmental timescales and diverse species, methods that reduce the need for task-specific retraining will facilitate more rapid integration of image acquisition with quantitative analysis. SAP therefore offers a practical framework for applying foundation-model segmentation to plant research, enabling reproducible and transferable workflows across experimental systems.

## Declarations

## List of abbreviations

Segment: SAM2 Anything Model 2
SAP: Segment Any Plant
IoU: Intersection over Union;

## Code Availability

The code is available at https://github.com/merozlab/plant-segmentation-app.

## Data Availability

The datasets generated and analyzed during this study are available on Zenodo at https://doi.org/10.5281/zenodo.18732705. This includes raw images and segmentation masks for the sunflower gravitropism and *Arabidopsis* root growth experiments, SAP-generated masks for the Lee et al. (9) and Strauss et al. (13) datasets, centerline validation data, and supplementary videos.

## Funding

Y.M. acknowledges support from the Israel Science Foundation Research Grant (ISF) no. 2307/22, and ERC grant GROWsmart 101165101. A.A. acknowledges support from the Lautman Foundation for his studies.

## Competing Interests

The author declares that they have no competing interests.

## Author’s Contributions

A.A.: conception, development of methodology; A.A. and Y.M.: formal analysis, conceptualization and investigation, visualization, writing original draft, review, and editing; Y.M. funding acquisition, supervision, and resources.

## ACKNOWLEDGEMENTS

The authors thank Águeda De la Vega for supplying *Arabidopsis* root imagery.

## Supplementary Information: Segment Any Plant (SAP)

### Supplementary Note A: Centerline Extraction Algorithm

The centerline extraction algorithm takes a binary segmentation mask of an elongated structure (e.g. a stem or root) and returns an ordered sequence of points along its medial axis. The method proceeds in five stages.

#### 1. Adaptive parameter estimation

Given the largest external contour of the mask, its points are redistributed uniformly by arc-length and analyzed with PCA. The minor principal axis is intersected with the mask boundary to estimate the structure’s width *w*. The length is estimated as *ℓ* = (*P* − 2*w*)*/*2, where *P* is the contour perimeter. Two parameters are then derived: the *edge fraction f* = (*w* + *ℓ/*2)*/*(2*w* + 2*ℓ*), which controls how much of the contour near each tip is used for endpoint localization, and the number of output centerline points *n* = ⌈*ℓ*⌉. Both parameters can also be set manually.

#### 2. Endpoint detection via skeletonization

The binary mask is thinned to a one-pixel-wide skeleton (18). Skeleton pixels with exactly one 8-connected neighbor are identified as endpoints. When more than two endpoints exist, the pair with the greatest Euclidean separation is selected. If fewer than two endpoints are found, the two most distant skeleton pixels are used instead.

#### 3. Tip localization

For each skeleton endpoint, a neighborhood of contour points is extracted. The neighborhood size equals *f* times the total number of contour points, centered on the contour point nearest to the skeleton endpoint. A minimum-area rectangle is fitted to each neighborhood via PCA. A line connecting the midpoints of the rectangle’s two shorter sides (i.e. aligned with the structure’s width at the tip) is extended and intersected with the contour. The intersection closest to the skeleton endpoint is taken as the *tip point*—the point on the contour that marks the true end of the structure.

#### 4. Contour splitting and averaging

The two tip points divide the contour into two open curves, corresponding to the two long edges of the elongated structure. Each curve is resampled to *n* uniformly spaced points by arc-length interpolation. The second curve is reversed so that corresponding points on the two edges face each other. The centerline is computed as the pointwise mean of the two resampled edge curves:

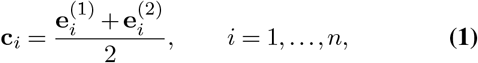

where 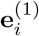 and 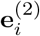 are corresponding points on the first and second edge curves, respectively.

#### 5. Validation and redistribution

When the structure has concave edges, the pointwise average of opposing contour points can fall outside the mask. Any such centerline point is discarded. The surviving points are then redistributed uniformly by arc-length interpolation to restore a regularly spaced *n*-point centerline. The output is ordered so that the first point has the smaller *x*-coordinate.

By averaging opposite contour edges rather than relying solely on morphological thinning, this approach produces a centerline that is equidistant from both long boundaries of the structure, even at the tips where classical skeletonization methods tend to deviate toward corners.

## Supplementary Videos

**Supplementary Video S1**. Video demonstration of the SAP workflow, corresponding to Fig. 1. **Supplementary Video S2**. SAP segmentation of *Arabidopsis thaliana* rosette growth over 9 days (9). **Supplementary Video S3**. SAP segmentation and centerline extraction of sunflower (*Helianthus annuus*) gravitropism. **Supplementary Video S4**. SAP segmentation and centerline extraction of *Arabidopsis thaliana* root growth. **Supplementary Video S5**. SAP segmentation of individual *Arabidopsis* root cells in confocal *z*-stack imagery (13). **Supplementary Video S6**. Cell-level segmentation of a growing *Arabidopsis* root in confocal time-lapse imagery (14), demonstrating SAP’s ability to track individual cells across frames.

**Fig. 2. Supplementary Video S2.**
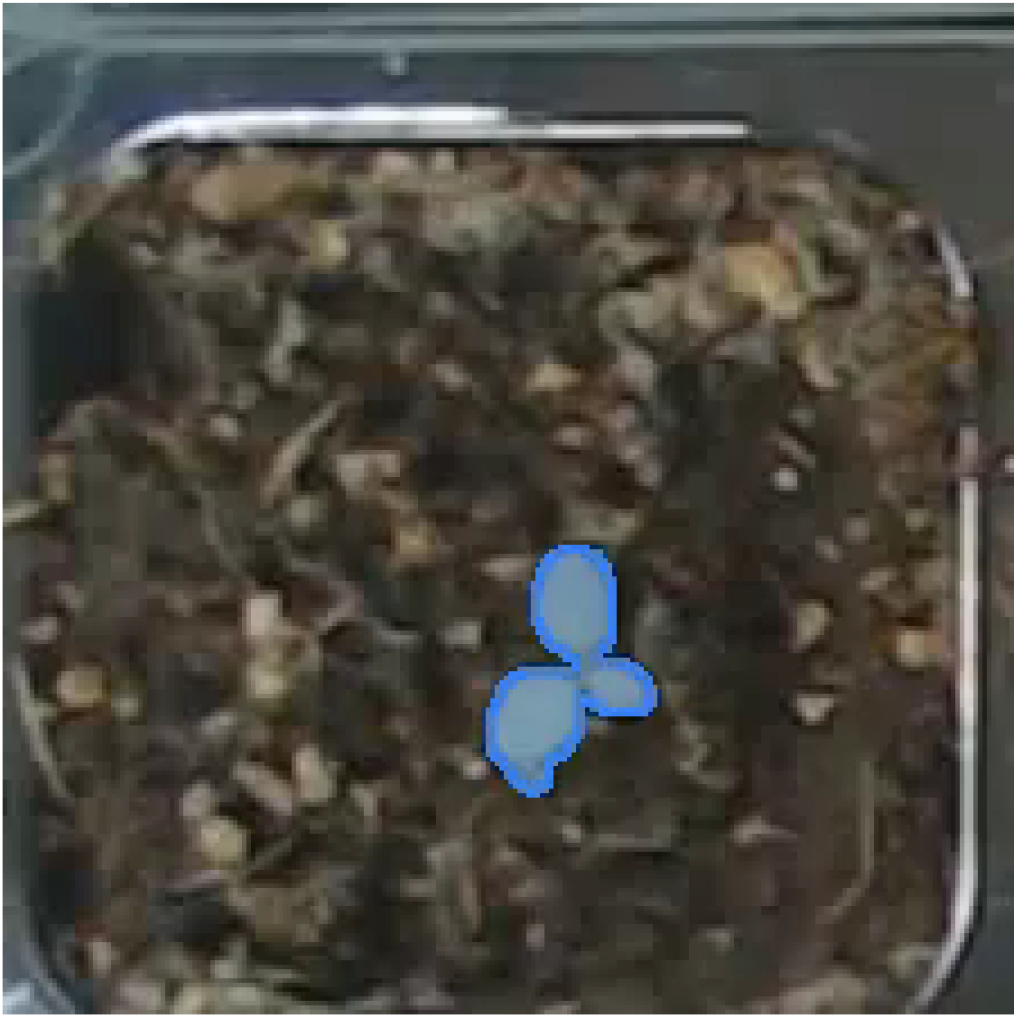
SAP segmentation of *Arabidopsis thaliana* rosette growth over 9 days (9). Top-down view of a rosette in a soil pot; SAP masks (blue) delineate leaf area from a single-frame prompt, propagated across a 9-day growth series.

**Fig. 3. Supplementary Video S3.**
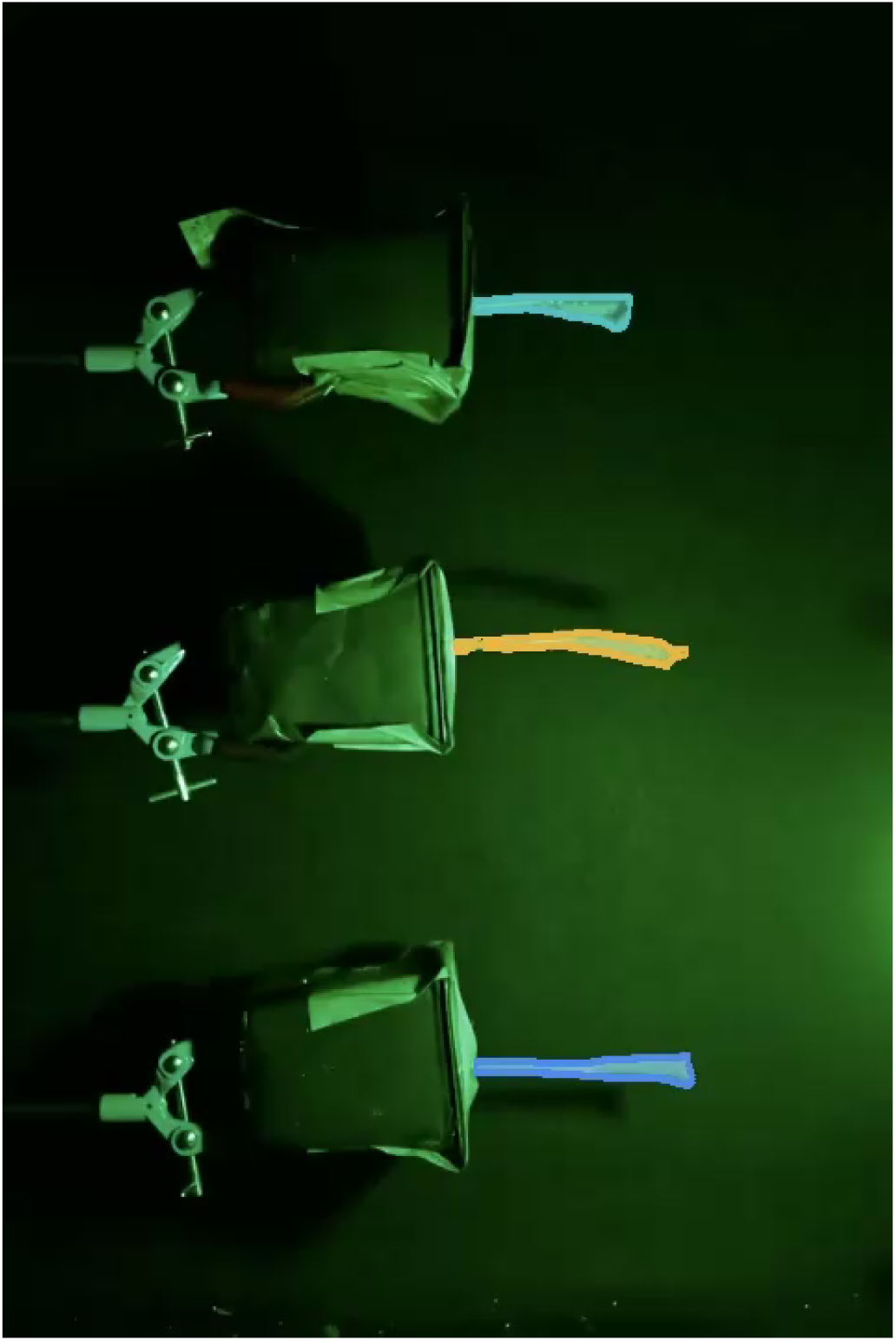
SAP segmentation and centerline extraction of sunflower (*Helianthus annuus*) gravitropism. Three defoliated sunflower stems at the onset of a gravitropism experiment; SAP masks and extracted centerlines track stem reorientation over 32 hours.

**Fig. 4. Supplementary Video S4.**
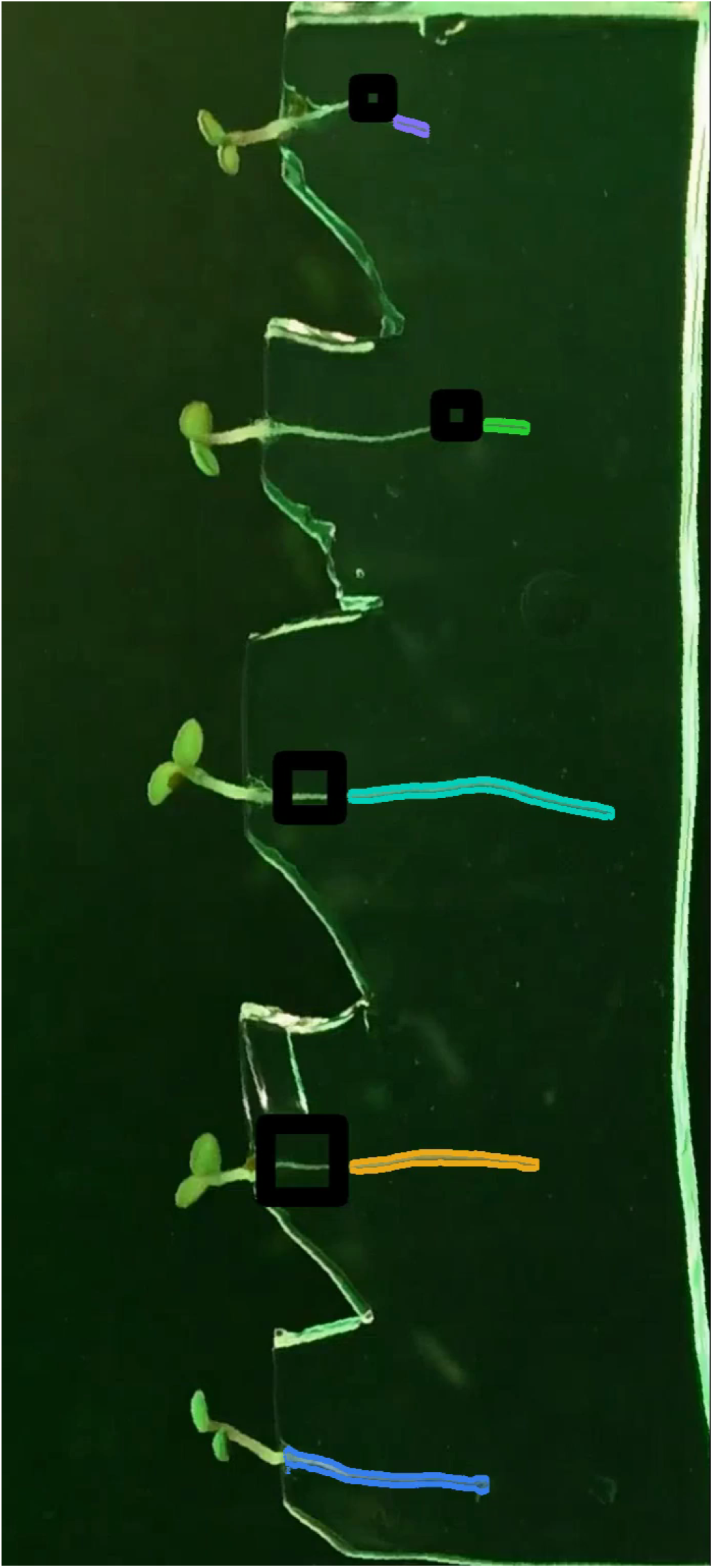
SAP segmentation and centerline extraction of *Arabidopsis thaliana* root growth. Seedlings growing on a vertical plate under dim green light; SAP segmentation masks and centerlines (colored by plant) enable root elongation analysis over 30 hours.

**Fig. 5. Supplementary Video S5.**
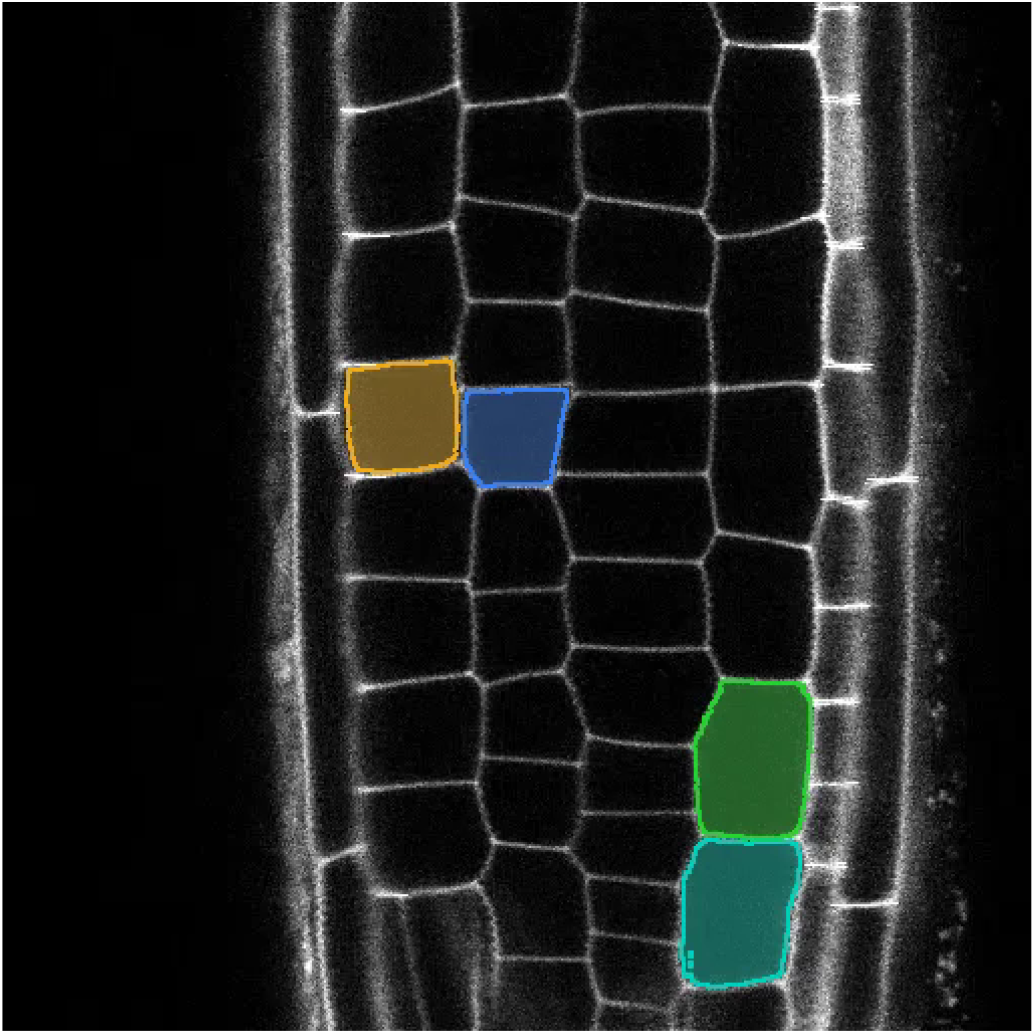
SAP segmentation of individual *Arabidopsis* root cells in confocal *z*-stack imagery (13). Confocal *z*-slice of an *Arabidopsis* root stained with Propidium Iodide; four individual cells are segmented (yellow, blue, teal, green) and tracked through a 51-slice subvolume.

**Fig. 6. Supplementary Video S6.**
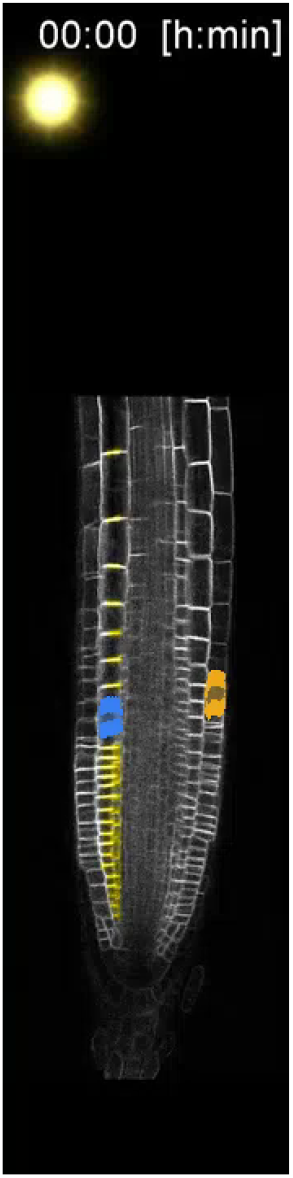
Cell-level segmentation of a growing *Arabidopsis* root in confocal time-lapse imagery (14), demonstrating SAP’s ability to track individual cells across frames. Confocal time-lapse of a growing *Arabidopsis* root tip with cell-level SAP segmentation across successive time points.

## Notes

https://doi.org/10.5281/zenodo.18732705

### Competing Interest Statement

The authors have declared no competing interest.

### Summary of Updates

The manuscript was revised to strengthen the presentation of validation results and broaden the paper's scope to include spatial (z-stack) analysis alongside temporal sequences. The main changes include: adding a dataset summary table for quick reference, expanding Figure 2 to show validation across all three ground-truth datasets (adding sunflower), extending centerline validation to cover both the sunflower and Arabidopsis root datasets, and rewriting the results section to emphasize extraction of biologically meaningful quantities like growth rates and tropic responses. The supplementary figures were reorganized from a single composite into individual figures with proper cross-references. The abstract and discussion were updated to reflect the broader spatial+temporal scope, and the centerline extraction method description was expanded to highlight its novelty. Minor text and formatting fixes were applied throughout.

## Bibliography

1. Malia A Gehan, Noah Fahlgren, Arash Abbasi, Jeffrey C Berry, Steven T Callen, Leonardo Chavez, Andrew N Doust, Max J Feldman, Kerrigan B Gilbert, John G Hodge, et al. Plantcv v2: Image analysis software for high-throughput plant phenotyping. PeerJ, 5:e4088, 2017. doi:10.7717/peerj.4088.

2. Keiichi Mochida, Satoru Koda, Komaki Inoue, Takashi Hirayama, Shojiro Tanaka, Ryuei Nishii, and Farid Melgani. Computer vision-based phenotyping for improvement of plant productivity: a machine learning perspective. GigaScience, 8(1):giy153, 2019. doi:10.1093/gigascience/giy153.

3. Félix P. Hartmann, Hugo Chauvet-Thiry, Jérôme Franchel, Stéphane Ploquin, Bruno Moulia, Nathalie Leblanc-Fournier, and Mélanie Decourteix. Methods for a Quantitative Comparison of Gravitropism and Posture Control Over a Wide Range of Herbaceous and Woody Species, pages 117–131. Springer US, October 2021. ISBN 9781071616772. doi:10.1007/978-1-0716-1677-2_9.

4. Gabriella E. C. Gall, Talmo D. Pereira, Alex Jordan, and Yasmine Meroz. Fast estimation of plant growth dynamics using deep neural networks. Plant Methods, 18(1), February 2022. ISSN 1746-4811. doi:10.1186/s13007-022-00851-9.

5. Renaud Bastien, David Legland, Marjolaine Martin, Lucien Fregosi, Alexis Peaucelle, Stéphane Douady, Bruno Moulia, and Herman Höfte. Kymorod: a method for automated kinematic analysis of rod-shaped plant organs. The Plant Journal, 88(3):468–475, 2016. ISSN 1365-313X. doi:10.1111/tpj.13255.

6. Nicolás Gaggion, Federico Ariel, Vladimir Daric, Éric Lambert, Simon Legendre, Thomas Roulé, Alejandra Camoirano, Diego H Milone, Martin Crespi, Thomas Blein, and Enzo Ferrante. Chronoroot: High-throughput phenotyping by deep segmentation networks reveals novel temporal parameters of plant root system architecture. GigaScience, 10(7), July 2021. ISSN 2047-217X. doi:10.1093/gigascience/giab052.

7. Michael P. Pound, Jonathan A. Atkinson, Alexandra J. Townsend, Michael H. Wilson, Marcus Griffiths, Aaron S. Jackson, Adrian Bulat, Georgios Tzimiropoulos, Darren M. Wells, Erik H. Murchie, Tony P. Pridmore, and Andrew P. French. Deep machine learning provides state-of-the-art performance in image-based plant phenotyping. GigaScience, 6(10):gix083, 08 2017. ISSN 2047-217X. doi:10.1093/gigascience/gix083.

8. Chen Shen, Liantao Liu, Lingxiao Zhu, Jia Kang, Nan Wang, and Limin Shao. High-throughput in situ root image segmentation based on the improved deeplabv3+ method. Frontiers in Plant Science, 11, October 2020. ISSN 1664-462X. doi:10.3389/fpls.2020.576791.

9. Unseok Lee, Sungyul Chang, Gian Anantrio Putra, Hyoungseok Kim, and Dong Hwan Kim. An automated, high-throughput plant phenotyping system using machine learning-based plant segmentation and image analysis. PLOS ONE, 13(4):e0196615, April 2018. ISSN 1932-6203. doi:10.1371/journal.pone.0196615.

10. Yixiang Mao, Hejian Liu, Yao Wang, and Eric D. Brenner. A deep learning approach to track arabidopsis seedlings’ circumnutation from time-lapse videos. Plant Methods, 19(1), February 2023. ISSN 1746-4811. doi:10.1186/s13007-023-00984-5.

11. Nikhila Ravi, Valentin Gabeur, Yuan-Ting Hu, Ronghang Hu, Chaitanya Ryali, Tengyu Ma, Haitham Khedr, Roman Rädle, Chloe Rolland, Laura Gustafson, Eric Mintun, Junting Pan, Kalyan Vasudev Alwala, Nicolas Carion, Chao-Yuan Wu, Ross Girshick, Piotr Dollár, and Christoph Feichtenhofer. Sam 2: Segment anything in images and videos. arXiv preprint arXiv:2408.00714, 2024. doi:10.48550/ARXIV.2408.00714.

12. Lian Lei, Qiliang Yang, Ling Yang, Tao Shen, Ruoxi Wang, and Chengbiao Fu. Deep learning implementation of image segmentation in agricultural applications: a comprehensive review. Artificial Intelligence Review, 57(6), May 2024. ISSN 1573-7462. doi:10.1007/s10462-024-10775-6.

13. Sören Strauss, Adam Runions, Brendan Lane, Dennis Eschweiler, Namrata Bajpai, Nicola Trozzi, Anne-Lise Routier-Kierzkowska, Saiko Yoshida, Sylvia Rodrigues da Silveira, Athul Vijayan, Rachele Tofanelli, Mateusz Majda, Emillie Echevin, Constance Le Gloanec, Hana Bertrand-Rakusova, Milad Adibi, Kay Schneitz, George W. Bassel, Daniel Kierzkowski, Johannes Stegmaier, Miltos Tsiantis, and Richard S. Smith. Using positional information to provide context for biological image analysis with MorphoGraphX 2.0. eLife, 11:e72601, 2022. doi:10.7554/eLife.72601.

14. Daniel von Wangenheim, Robert Hauschild, Matyáš Fendrych, Vanessa Barone, Eva Benková, and Jiří Friml. Live tracking of moving samples in confocal microscopy for vertically grown roots. eLife, 6:e26792, jun 2017. ISSN 2050-084X. doi:10.7554/eLife.26792.

15. Md Atiqur Rahman and Yang Wang. Optimizing Intersection-Over-Union in Deep Neural Networks for Image Segmentation, pages 234–244. Springer International Publishing, 2016. ISBN 9783319508351. doi:10.1007/978-3-319-50835-1_22.

16. Zhifang Bi, Fumin Ma, Jiaxiong Guan, Jie Wu, Zhanli Liu, Yanwen Li, Fuzhong Li, and Juxia Li. Apple leaf disease severity grading based on deep learning and the drl-watershed algorithm. Scientific Reports, 15(30071), January 2025. ISSN 2045-2322. doi:10.1038/s41598-025-15246-8.

17. Kotaro T. Yamamoto and Ken Haga. Quantitative Measurements of Curvature Along the Growth Axis in Tropic Responses Using Free Software Environments, pages 223–234. Springer New York, New York, NY, 2019. ISBN 978-1-4939-9015-3. doi:10.1007/978-1-4939-9015-3_19.

18. T.C. Lee, R.L. Kashyap, and C.N. Chu. Building skeleton models via 3-d medial surface axis thinning algorithms. CVGIP: Graphical Models and Image Processing, 56(6):462–478, November 1994. ISSN 1049-9652. doi:10.1006/cgip.1994.1042.

19. Xiang Jiao, Huichun Zhang, Jiaqiang Zheng, Yue Yin, Guosu Wang, Ying Chen, Jun Yu, and Yufeng Ge. Comparative analysis of nonlinear growth curve models for arabidopsis thaliana rosette leaves. Acta Physiologiae Plantarum, 40(6), May 2018. ISSN 1861-1664. doi:10.1007/s11738-018-2686-8.

20. Renaud Bastien, Tomas Bohr, Bruno Moulia, and Stéphane Douady. Unifying model of shoot gravitropism reveals proprioception as a central feature of posture control in plants. Proceedings of the National Academy of Sciences, 110(2):755–760, December 2012. ISSN 1091-6490. doi:10.1073/pnas.1214301109.

